# Chill injury in human kidney tubule cells after subzero storage is not mitigated by antifreeze protein addition

**DOI:** 10.1101/2022.08.09.503393

**Authors:** Heather E. Tomalty, Laurie A. Graham, Virginia K. Walker, Peter L. Davies

## Abstract

By preventing freezing, antifreeze proteins (AFPs) can permit cells and organs to be stored at subzero temperatures. As metabolic rates decrease with decreasing temperature, subzero static cold storage (SCS) could provide more time for tissue matching and potentially lead to fewer discarded organs. Human kidneys are generally stored for under 24 h and the tubule epithelium is known to be particularly sensitive to SCS. Here, telomerase-immortalized proximal-tubule epithelial cells from humans, which closely resemble their progenitors, were used as a proxy to assess the potential benefit of subzero SCS for kidneys. The effects of hyperactive AFPs from a beetle and Cryostasis Storage Solution were compared to University of Wisconsin Solution at standard SCS temperatures (4 °C) and at −6 °C for up to six days. Although the AFPs helped guard against freezing, lower storage temperatures under these conditions were not beneficial. Compared to cells at 4 °C, those stored at −6 °C showed decreased viability as well as increased lactate dehydrogenase release and apoptosis. This suggests that this kidney cell type might be prone to chilling injury and that the addition of AFPs to enable subzero storage may not be effective for increasing storage times.

## Introduction

Considerable effort has been directed towards developing new approaches to improve the preservation of cells, tissues, and whole organs that are compromised when subjected to freezing and thawing during conventional cryopreservation. Static cold storage (SCS) at near zero temperatures is routinely used, but storage times are limited ^1^. For example, hearts can only be stored for 4–6 h ^2^, livers for around 12 h ^3^ and kidneys stored for under 12 h lead to better patient outcomes, while those stored for over 24 h have a high rate of failure ^4^. There are limited opportunities for tissue matching and/or transport of organs to recipients within these time frames, leading to organ wastage, long waiting lists, graft rejection, and patient deaths ^1-3^. Therefore, the ability to extend this time frame would be beneficial, easing the logistical demands faced by transplant communities.

Storing freeze-sensitive biological materials at subzero temperatures in an unfrozen state (subzero static cold storage (SZ-SCS)) could provide a simple, cost-effective solution as the metabolic processes that can lead to ischaemic damage decrease with decreasing temperature ^5^. The feasibility of this approach was best demonstrated using rat livers that were successfully transplanted after 4 days of storage at −6 °C ^6^. Despite this initial success, SZ-SCS does have some drawbacks, including chilling injury resulting when susceptible cells or tissues are placed at low temperatures ^7^, as well as the risk of ice formation. As previously reported, ice can be avoided through the use of colligative antifreeze agents; however, they must be used at high concentrations to be effective, and have varying toxicities that increase with concentration ^8^. For example, the rat liver study cited employed 5% polyethylene glycol (PEG) 35000, along with 0.2 M of a non-metabolizable glucose derivative that can cross the cell membrane to reduce cellular dehydration ^6^. A similar supercooling protocol was successfully applied to human livers cooled to −4 °C, with several modifications, including the addition of 10% glycerol, as their 200-fold greater size makes them more prone to ice nucleation. The authors also noted that nucleation could be induced by jostling if transport of the organ was required ^3^. Given this, SZ-SCS would benefit if less toxic antifreeze agents could be used.

Antifreeze proteins (AFPs) are promising agents as they depress the freezing point at much lower concentrations than the colligative antifreezes mentioned above (Table 1). They do so by binding irreversibly to the surface of ice crystals, making further growth thermodynamically unfavorable ^9,10^. The difference between the melting point and the non-equilibrium freezing point can be accurately measured and it is denoted as the thermal hysteresis (TH) activity ^11^. AFPs have been used in a variety of industrial and medical applications, to prevent freezing or to moderate the effects of ice formation. These applications include the preservation of organs, tissues, and cells in both the supercooled and the frozen state ^12^. AFPs have been found in a variety of organisms, but it is the AFPs of certain terrestrial insects that are of particular interest for SZ-SCS as they are classified as hyperactive, having >10-fold higher activity compared to moderately-active AFPs from fish ^13^. The antifreeze proteins of the yellow mealworm beetle, *Tenebrio molitor* (*Tm*AFP), were chosen for this study, as a synergistic mixture of isoforms ^14^ can be purified directly from the insects, using a simple ice-affinity procedure ^15^. They can provide 6 °C of freeze protection, which is unattainable using fish AFPs, but which can be achieved with glycerol, albeit at a 900-fold higher mass concentration (Table 1).

**Table 1.** Freezing point depression capabilities of various compounds.

| Compound | Concentration |  | Freezing Point Depression (°C) |
| --- | --- | --- | --- |
|  | Molarity | mg/mL |  |
| Glycerol <sup>42</sup> | 1.4 | 126 | -2.3 |
| Glycerol | 2.2 | 200 | -6 |
| 3-O-methyl-D-glucose <sup>43</sup> | 0.2 | 39 | -0.47 |
| Type II AFP <sup>44</sup> | $1.2 \times 10^{-3}$ | 17 | -0.46 |
| Type III AFP <sup>45</sup> | $7.2 \times 10^{-4}$ | 5 | -0.75 |
| *Tm*AFP <sup>20</sup> | $2.4 \times 10^{-5}$ | 0.22 | -6 |
| *Tm*AFP + CrS <sup>1</sup> | $1.1 \times 10^{-5}$ | 0.1 | -7.3 |

Tubular atrophy is a hallmark of graft dysfunction and kidney disease ^16,17^, with apoptosis of tubule epithelial cells being implicated ^18^. Therefore, the RPTEC/TERT1 cell line was chosen to investigate the response of human renal cells to SZ-SCS. These cells were derived from proximal-tubule epithelial cells by transformation with human telomerase and they retain both chromosomal and normal functional stability over many generations ^19^. Therefore, they are a better proxy for evaluating the ability of SZ-SCS to preserve human kidney function than cell lines derived from tumors, embryonic cells or cells from other species. In addition, small volumes of cells can be maintained at high subzero temperatures without freezing, enabling evaluation of the effect of the presence and absence of low concentrations of *Tm*AFP, sufficient to confer 6 °C of freezing protection, in combination with Cryostasis Solution™ (CrS), on cell preservation. Our result show that *Tm*AFP is not deleterious, but neither is it demonstrably protective at the concentrations used, that CrS solution improves survival during SZ-SCS at −6 °C relative to University of Wisconsin (UW) solution, and that SZ-SCS does not improve the long-term survival of RPTEC/TERT1 cells under the conditions tested.

## Materials and Methods

### TmAFP purification

*Tm*AFP was purified from homogenized cold-acclimated *T. molitor* larvae as previously described ^20^. As *Tm*AFP is a mixture of several isoforms that differ in size and/or charge, five rounds of rotary ice-affinity purification was used since this technique is highly selective for proteins that bind to ice, irrespective of their other properties ^15^. The final ice fraction was melted while NH_4_HCO_3_ was slowly added to a final concentration of 20 mM. The AFP was concentrated approximately 100-fold using an Amicon stirred cell and centrifugal filters (3,000 MWCO; Sigma-Aldrich, Oakville, Canada) and aliquots were snap-frozen and stored at −80 °C. Amino acid analysis was performed at SPARC BioCentre (The Hospital for Sick Children, Toronto, Canada) to determine final *Tm*AFP concentrations, which ranged from ∼2 to 4 mg/mL.

### Antifreeze activity

TH (thermal hysteresis) activity is the difference between an ice crystal’s melting point and its non-equilibrium freezing point, where the ice crystal shows sudden growth. It was measured using a nanolitre osmometer ^11^ with *Tm*AFP in either 20 mM NH_4_HCO_3_ or the proprietary Cryostasis Solution (CrS) (Cryostasis Ltd., Westport, Canada).

### Ice nucleation

The ability of *Tm*AFP to prevent ice nucleation was assessed using 1 mL aliquots of either CrS or CrS + *Tm*AFP cell storage medium in the presence or absence of ∼200 ng of lyophilized outer membranes of *Pseudomonas syringae* that have extremely active ice nucleation proteins (INPs) (Ward’s Natural Science, Rochester, USA). Samples were held at either -4, -5, or -6 °C for 18 h and the experiment was repeated three times with triplicates, for a total of nine vials per treatment.

### Cell culture growth conditions

The human renal proximal tubule cell line, RPTEC/TERT1 (ATCC CRL-4031), was obtained from Cedarlane (Burlington, Canada) and grown according to ATCC guidelines. Cells were cultured in Dulbecco Modified Eagle Medium - Ham’s F-12 supplemented with ATCC hTERT Immortalized RPTEC Growth Kit (Cedarlane) and 100 μg/mL G418 antibiotic (Sigma-Aldrich) under standard conditions (37 °C, 5% CO_2_), hereafter abbreviated as DMEM. Cells were subcultured once confluent at either a 1:3 or 1:4 ratio. All experiments were performed using cells between passages 4 to 10.

### Supercooling experiments

To investigate the efficacy of supercooled preservation, three preservation solutions were tested: University of Wisconsin (UW), CrS, and CrS + *Tm*AFP. RPTEC/TERT1 cells were grown to confluence and detached using 0.25% trypsin in 0.5 mM EDTA, after which 0.1% soybean trypsin inhibitor (Sigma-Aldrich) was added. Cell counts were obtained using a Moxi Z mini cell counter (VWR, Mississauga, Canada) and aliquots of 1 × 10^6^ cells were placed into 2-mL cryovials (Thermo Fisher Scientific, Waltham, USA). Cells were pelleted by centrifugation for 5 min at 160 x g, washed with 1 mL of phosphate-buffered saline (PBS) pH 7.4, pelleted again as above, then resuspended in 150 μL of preservation solution. Hypoxic conditions were generated by overlaying 350 μL of sterile, heavy mineral oil (Sigma-Aldrich) to cells in preservation solution that had been pelleted as above. Vials were stored at three temperatures (37 °C, 4 °C, −6 °C) for three different durations (1, 3, or 6 days), after which the mineral oil was removed and 0.5 mL of 21 °C cell medium was added to each vial. Cell pellets were gently resuspended, and a portion was assessed immediately for viability (see below). The balance of the cells were incubated at 37 °C for 1 h to begin recovery, after which the cells were centrifuged as above and resuspended in 1 mL of cell medium. Cells were plated according to the protocols outlined below.

### Viability counts

Cell viability counts were obtained immediately following supercooling and after a 20-h recovery period. Cells (0.3 × 10^6^) were resuspended in PBS and mixed 1:1 with 0.4% trypan blue (Sigma-Aldrich) and counts were manually determined using a hemocytometer. For the 20-h recovery counts, cells were plated at 0.3 × 10^6^ cells per well in 6-well plates and incubated under standard growth conditions (37 °C, 5% CO_2_) prior to viability assessment. Percent viability was calculated as the number of live cells divided by the total number of cells.

### Lactate dehydrogenase (LDH) release

Cells were plated in a 96-well plate at a density of 1 × 10^5^ cells per well and LDH release was assessed after a 20-h recovery period as an indirect measurement of necrotic cell death. LDH was measured using the Pierce™ LDH Cytotoxicity Assay Kit (Thermo Fisher Scientific) according to manufacturer’s directions. Absorbance of each well was determined at 490 nm, followed by a background reading at 680 nm using a SpectraMax Gemini plate reader (Molecular Devices, San Jose, USA). Background readings were subtracted from the 490 nm readings and LDH release calculated as a percentage of total LDH activity present within control wells containing lysed cells.

### TUNEL staining

After supercooling, cells were plated in a 96-well plate at a density of 2 × 10^5^ cells per well and held at 37 °C, 5 % CO_2_ for 24 h. Apoptotic cells were assessed using a Roche *in Situ* Cell Death Detection Kit, Fluorescein (Sigma-Aldrich) according to manufacturer’s directions. The cells were also stained using 300 nM DAPI solution (Thermo Fisher Scientific). Fluorescence images were captured on an Olympus IX83 inverted fluorescence microscope, with a minimum of five images captured for each well. Image software ^21^ was used to count fluorescent nuclei and the percent of apoptotic cells was calculated by dividing the number of TUNEL positive nuclei by the total number of nuclei present in each field of view.

### Long-term growth

Cells were seeded at 3 × 10^6^ cells per well on a 6-well plate and growth was monitored for 10 days or until the cells reached confluence, which ever came first. Images of the cells were taken every 3 days using an Olympus IX83 microscope (five images per well). The percent confluence for each image was calculated using the PHANTAST plug-in for FIJI ^22,23^.

### Statistical analysis

Data were analyzed using GraphPad Prism software (GraphPad Software, San Diego, USA). All results are reported as means ± standard error of the mean (SEM) and p<0.05 was considered statistically significant.

## Results

### TmAFP is highly effective for freeze inhibition

*Tm*AFP purified from insects is a mixture of isoforms that confers around 4 °C of freeze protection at a modest concentration of only 100 μg/mL in buffer (Fig. 1). However, it is more effective when used with CrS solution for two reasons. First, the thermal hysteresis (TH) activity is increased about 1 °C relative to AFP in buffer alone, to over 5 °C (Fig. 1). Second, CrS colligatively lowers the freezing point by 2 °C ^24^. Therefore, CrS + 100 μg/mL of *Tm*AFP provides 7.3 °C of combined protection, at a mass concentration three magnitudes lower than would be needed using glycerol (Table 1), ensuring that freezing will not occur during SZ-SCS at −6 °C.

**Figure 1.**
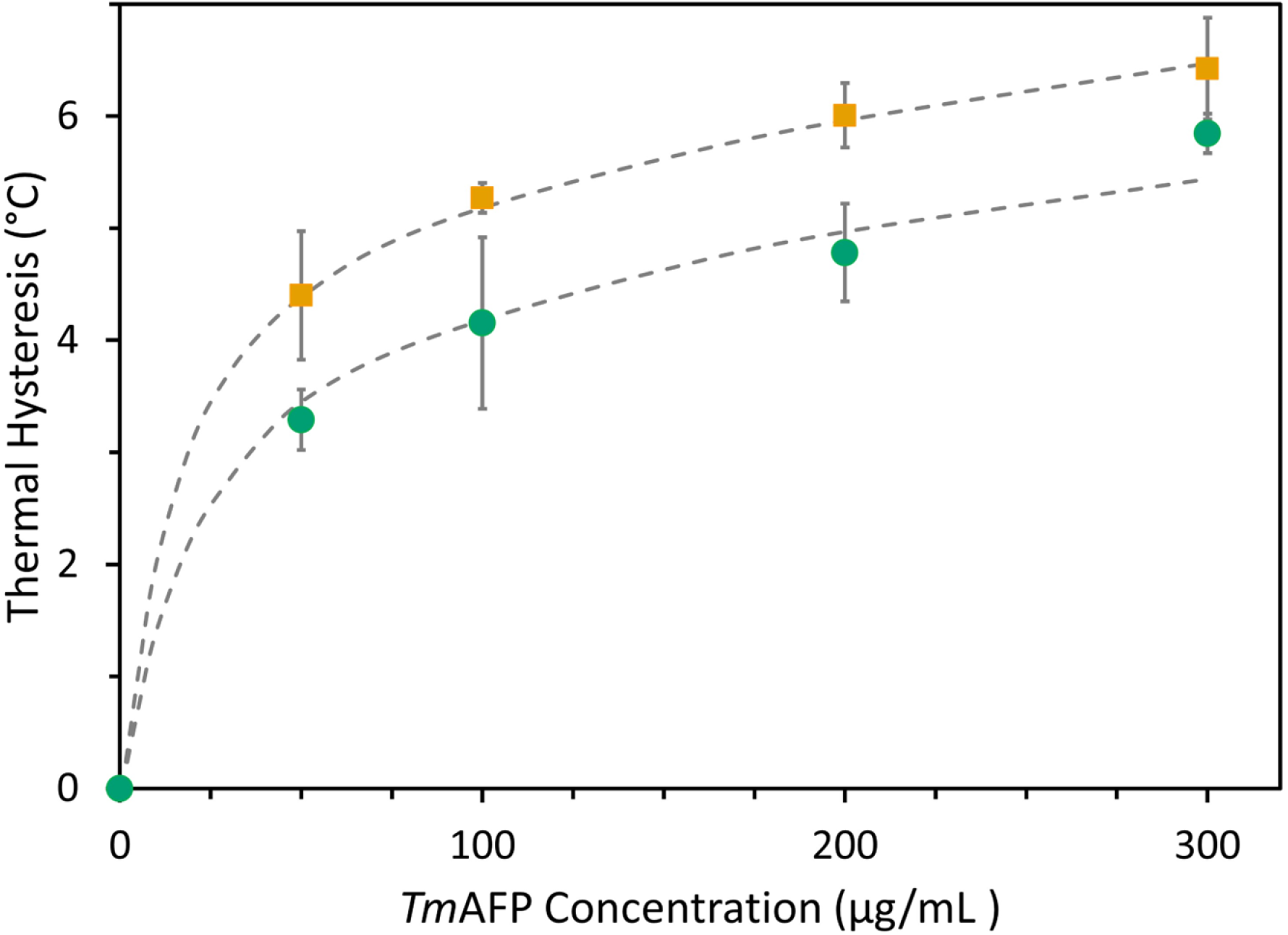
TH activity of *Tm*AFP at various concentrations in 20 mM ammonium bicarbonate (green circles, lower plot) or CrS solution (orange squares, upper plot). Error bars indicate standard deviation for three or more readings. TH was significantly different at each concentration (excepting zero) between the two solutions (t-tests, p < 0.05).

The effect of freezing damage was assessed in RPTEC/TERT1 cells by the addition of a potent ice nucleation protein (INPs), in the absence of AFP, during storage at -6 °C. All samples froze and the viability dropped to close to zero, underscoring the importance of eliminating freezing (Fig. 2). To evaluate whether this concentration of *Tm*AFP was likely to be effective at preventing heterogeneous ice nucleation in large organs that are prone to freezing at high subzero temperatures ^3^, INP was added to CrS or CrS + *Tm*AFP solutions (Table 2A). The volume used (1 mL) is around three orders of magnitude greater than the 1 – 2 μL typically used in nucleation assays ^25^. In the absence of *Tm*AFP, INP induced freezing in all tubes at −5 °C, which is only 3 °C below the normal freezing temperature of this solution. When *Tm*AFP was present, freezing was completely prevented at −5 °C, and only one of nine vials froze at −6 °C (Table 2).

**Table 2.** Suppression of an active ice nucleator (200 ng) by 100 μg/mL *Tm*AFP in 1 mL samples held for 18 h at three different temperatures.

| Solution | % Frozen at each holding temperature |  |  |
| --- | --- | --- | --- |
|  | -4 °C | -5 °C | -6 °C |
| CrS | 0 | 0 | 0 |
| CrS + INP | 0 | 100 | 100 |
| CrS + *Tm*AFP | 0 | 0 | 0 |
| CrS + *Tm*AFP + INP | 0 | 0 | 11 |

**Figure 2.**
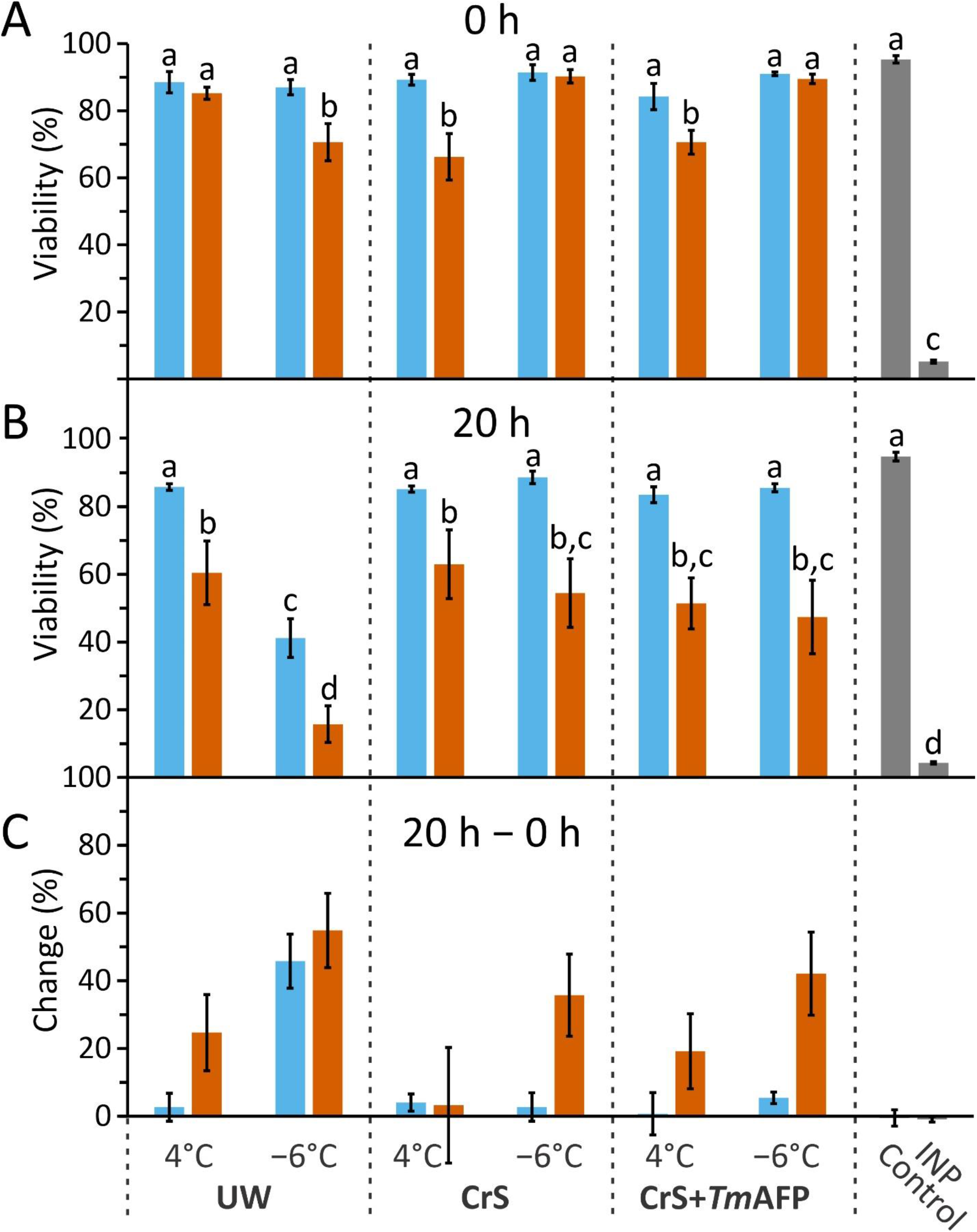
Viability of RPTEC/TERT1 cells after storage for three days (sky blue) or six days (reddish orange) in three different storage solutions ((UW, CrS, and CrS + *Tm*AFP) after conventional SCS (4 °C) or SZ-SCS (−6 °C). A) Viability immediately after cold storage. B) Viability after 20 h of recovery at 37 °C following cold storage. C) Change in cell viability during the 20 h recovery phase (difference between viability at 0 h and viability at 20 h). Cells from an actively growing culture (control, >95% viable) and cells that were frozen in CrS at -6 °C (<5% viable) by addition of ice nucleating protein (INP) are in grey. Error bars show the standard deviation of the five replicates. The letters (a to d) represent statistical differences determined by three-way ANOVA analysis with the same letter indicating that the sample means are not significantly different from each other.

## Cell death in RPTEC/TERT1 cells during cold storage is low in CrS solution at subzero temperatures

RPTEC/TERT1 cells were stored under oil for 1, 3 or 6 days, to mimic the hypoxia encountered by organs during SCS. In the absence of cooling, 29%–42% of the cells had died after only 1 day, 65%–88% after three days, and by day 6, almost all the cells were dead (Supplementary Figure 1). This occurred irrespective of whether nutritive medium or storage medium was used, indicating that the cells were compromised.

To assess the differences between SCS and SZ-SCS, cells were incubated at 4 °C or −6 °C in three different storage solutions (UW, CrS, CrS + *Tm*AFP) for one, three or six days. Cell viability was assessed by trypan blue staining immediately after the cells were removed from storage, to estimate cell death in storage. After one day, viability was similarly high (>88%) in both the 4 °C and −6 °C groups, so these data are presented separately in Supplementary Figure 2. The only exception was the UW sample stored at −6 °C, where viability decreased to 81%, which suggests that CrS solution was superior to UW for short-term SZ-SCS.

When cells were stored for three days at 4 °C, viabilities exceeded 84% and there was no significant difference between storage solutions (Fig. 2A, sky blue). However, when they were stored for six days at 4 °C (Fig. 2A, reddish orange), the cells in CrS ± *Tm*AFP showed a modest but statistically significant decrease in viability to around 70%, whereas the viability in UW remained high. When the storage temperature was −6 °C, this trend was reversed, with cells in UW solution having the lowest viability (71%) and those in CrS having viabilities of at least 90%. Therefore, CrS solution, with or without *Tm*AFP, appears beneficial for storage at −6 ° and the viability remained high even after six days, which initially suggested that SZ-SCS could be beneficial for extending storage time.

## Longer storage results in more cell death in cells recovering from SZ-SCS

The ability of cells to recover after storage at either 4 °C or −6 °C was assessed using trypan blue after 20 h of incubation at 37 °C in nutritive medium. The 1-day samples are shown in Supplementary Fig. 2 as all (except for UW at −6 °C, which showed only 76% viability) were over 90% viable. After three days of storage, samples stored at either temperature were over 83% viable (Fig. 2B, sky blue).

Additional death was minimal during recovery of three-day stored cells (Fig. 2C), with the one exception that cells stored in UW solution at −6 °C showed a 46% decrease in viability during recovery (Fig. 2C), again indicating that CrS was superior to UW for SZ-SCS of these cells over three days.

When the cells were subjected to 6 days of storage, they showed a significant drop in viability during the recovery period. This ranged from 47% to 63% (Fig. 2B,C, reddish orange), except with UW solution at −6 °C, where the viability dropped to only 16%. The differences between the viability of cells stored at 4 ° or −6 °C after recovery, at either the three-day or six-day timepoint, were not significant (Fig. 2B), despite the apparent benefit observed prior to recovery of cells stored at −6 °C in CrS solution (Fig. 2A). This can be attributed to greater cell death during recovery in the CrS-stored samples (Fig. 2C). This suggests that the CrS solution only delayed cell death at −6 °C, with an equivalent number of cells succumbing to chilling injury relative to those stored at 4 °C after the recovery period. Therefore, there is no apparent benefit to SZ-SCS for this cell type in these storage solutions. One notable observation is that there is no significant difference between storage in CrS or CrS supplemented with *Tm*AFP. This suggests that the AFP neither protects nor damages the cells at the low concentrations used here.

### Membrane leakage is highest in cells stored in UW solution

Lactate dehydrogenase (LDH) released into the medium during the 20 h recovery from storage conditions indicates damage to the plasma membrane and cytosolic protein leakage. LDH activity was significantly higher in cells stored in UW solution at −6 °C, with 22% and 28% LDH release over cells subjected to three days and six days of cold storage, respectively (Figure 3). The same samples showed 46% and 55% increases in cell death, respectively, during storage (Figure 2C). LDH release was lower in cells stored in CrS solution ± *Tm*AFP, and there was no statistically significant difference between the two storage temperatures, which is consistent with the conclusion that SZ-SCS is not beneficial for this cell type under these conditions.

**Figure 3.**
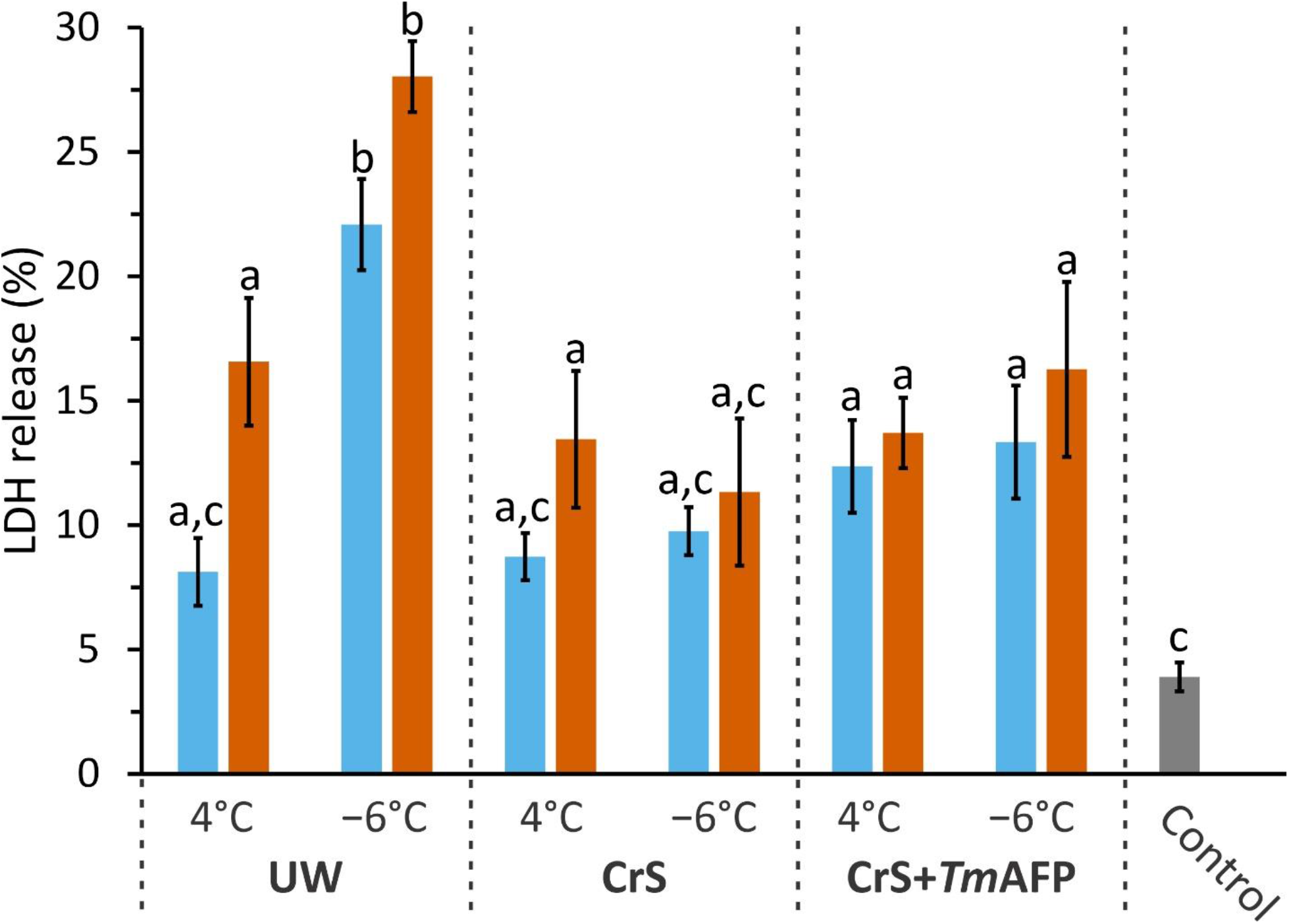
LDH release during the 20-h recovery period at 37 °C, for samples treated and colored as in Fig. 2. The error bars represent the standard deviation of triplicates. The percentage is relative to the LDH release from fully lysed cells (control, grey). Statistics are as described in Fig. 2.

### Apoptosis is a significant cause of cell death after cold storage

The TUNEL assay was used to assess late-stage apoptosis via fluorescent labelling of DNA ends generated during DNA fragmentation. Apoptotic cell death was below 5% in cells stored at 4 °C for three days, irrespective of storage solution (Fig. 4), and was not significantly different than the negative control. Cell death was noticeably higher following SZ-SCS in UW solution, with 25% of the cells classified as apoptotic after storage for only three days. After six days of storage at −6 °C, DNA fragmentation was observed in over 20% of the cells in all storage solutions. These data indicate that apoptosis is a significant contributor to the loss of cell viability that occurs during SZ-SCS relative to SCS, particularly during extended storage for six days, and this is evident in images of cells stained with both DAPI and TUNEL (Fig. 5).

**Figure 4.**
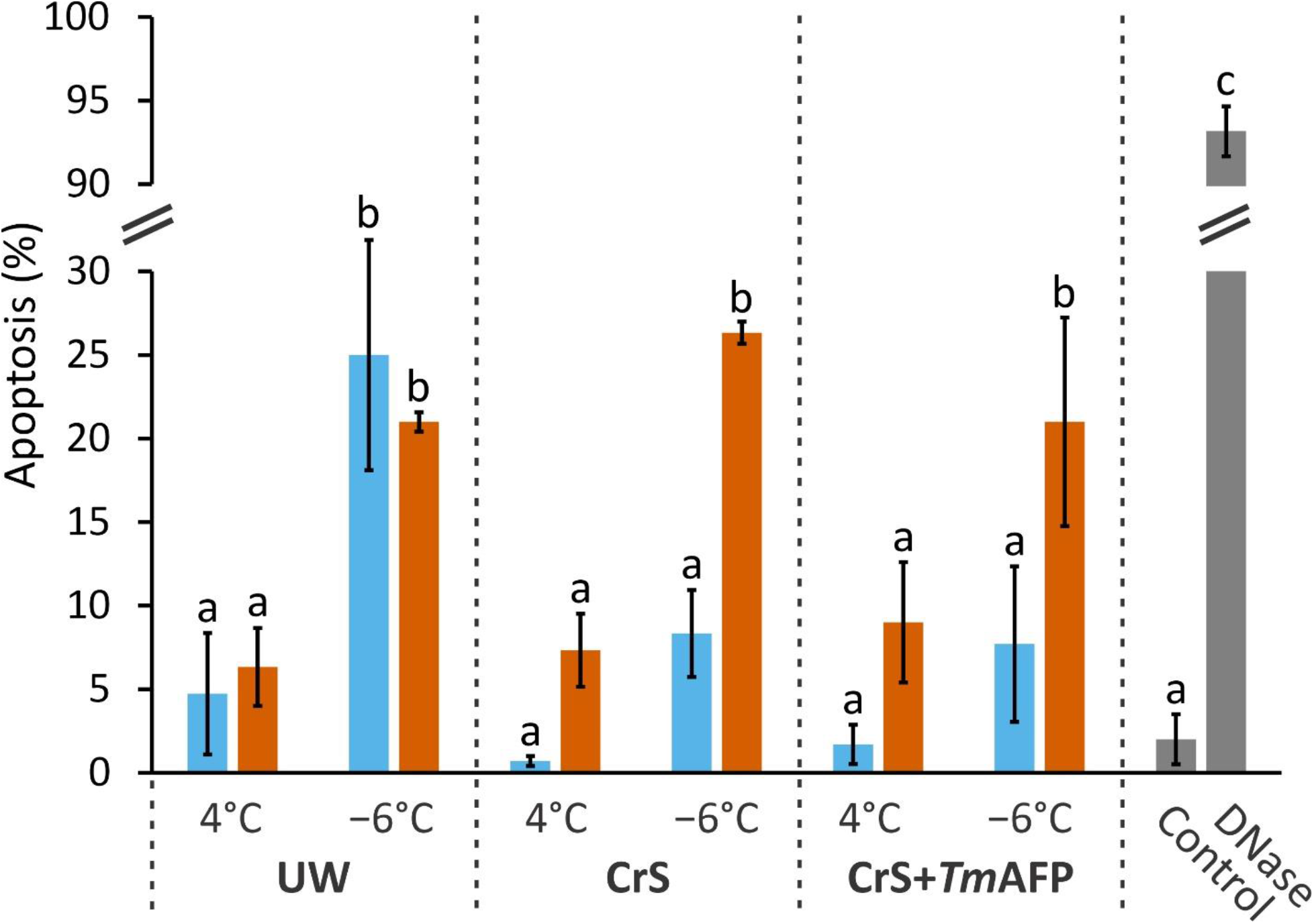
Apoptotic cells detected using the TUNEL assay on cells following a 24-h recovery period at 37 °C, for samples treated and colored as in Fig. 2. The apoptosis percentage is the number of TUNEL positive cells relative to the total number of cells stained by DAPI. The error bars represent the standard deviation of triplicates. Actively-growing cultures were used for the negative control (control, grey) and the positive control (DNase treated, grey). Statistics are as described in Fig. 2.

**Figure 5.**
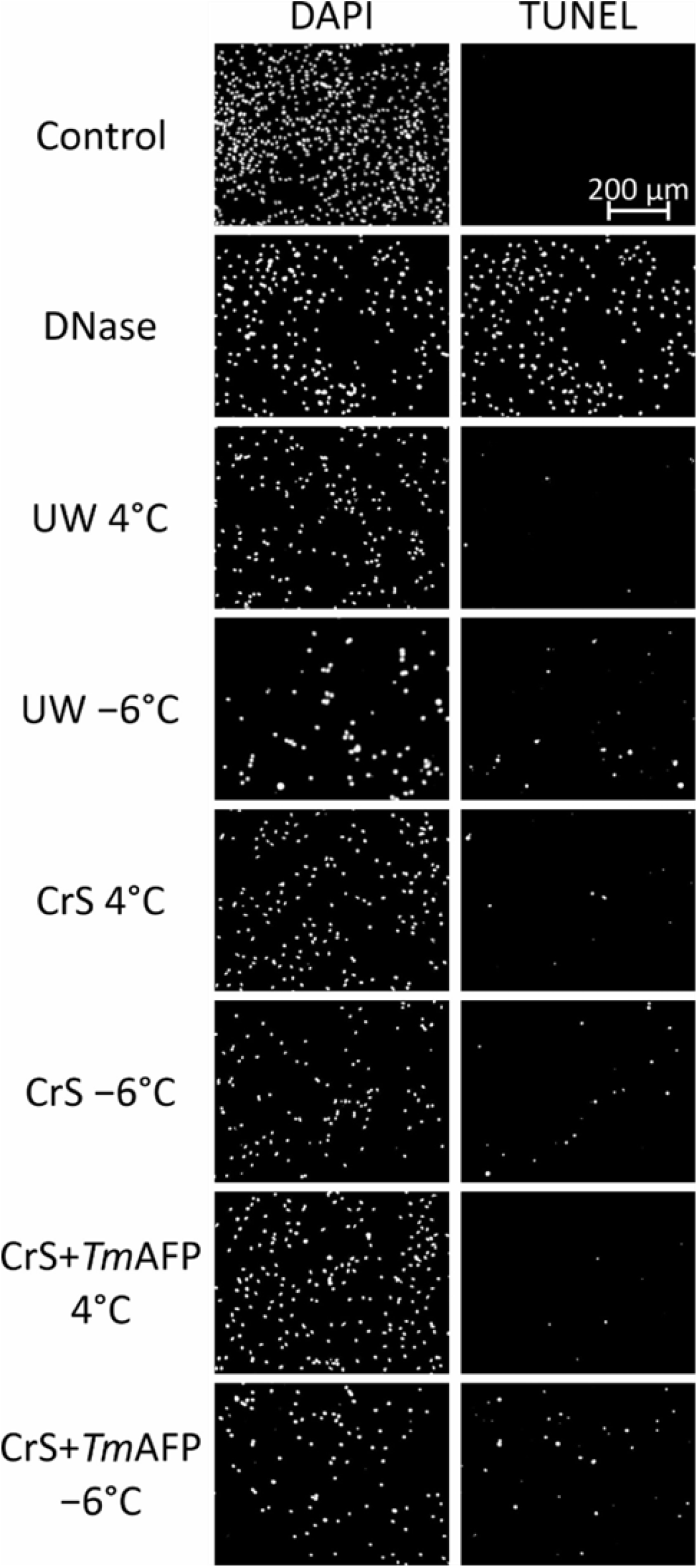
Representative images of DAPI-(left) and TUNEL-stained cells (right) from a subset of the samples used to generate Fig. 4. All samples (except controls) were stored at the indicated temperatures for six days. Images were converted to greyscale for ease of viewing.

One consistent difference between the SZ-SCS samples and those stored at the warmer temperature was that there was a notable decrease in cell density by DAPI staining. This suggests that a portion of the cells failed to adhere to the plate during the recovery period. This discrepancy is discussed further below. The lack of any significant differences between cells stored in CrS in the presence or absence of *Tm*AFP indicates that the AFP neither induces nor prevents apoptosis at the concentration used.

### Subzero storage is detrimental to the long-term growth of RPTEC/TERT1 cells

Growth of the cells following cold storage was evaluated by plating equal numbers of cells and monitoring growth over ten days (Fig. 6). By day seven, both control cells (black dotted line) and those stored at 4 °C for one day reached full confluence (Fig. 6A). In contrast, those stored at −6 °C for one day, in all storage solutions (Fig. 6A, dashed lines), showed a significant growth delay with ≤60% confluency, despite showing little loss of viability by trypan blue staining (Supplementary Fig. 2). This suggests the cells experience some chilling injury during SZ-SCS that inhibits proliferation.

**Figure 6.**
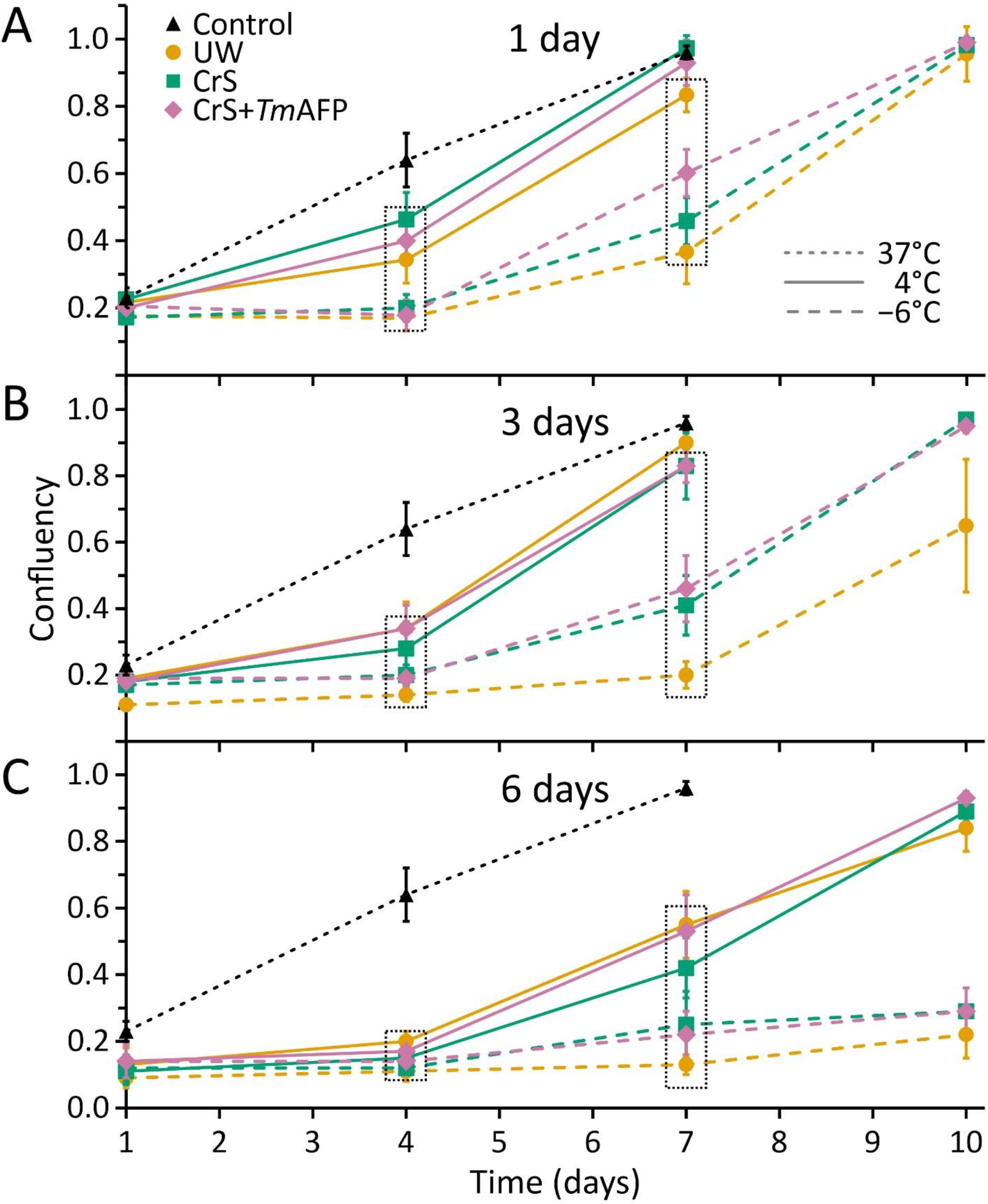
Recovery and long-term growth of cells following SCS and SZ-SCS. Cells were stored in UW solution (orange with circular markers), CrS solution (green with square markers) or CrS + *Tm*AFP (purple with diamond markers) at 4 °C (solid lines) or −6 °C (dashed lines). Control cells (no SCS) are shown in black with triangular markers. A) Percentage of surface occupied by cells (confluency) that were stored for one day and then plated and allowed to proliferate for ten days. B) Confluency of cells stored for three days. C) Confluency of cells stored for six days. Dashed boxes encompass confluencies at each time point that are significantly different from the control, as determined by unpaired t-tests.

An increase in storage time to three days resulted in slightly slower rates of growth, again with the cells stored at −6 °C showing the greatest delays. One notable difference is that the cells in UW solution at the lower temperature show an additional growth lag of approximately two days. However, after six days of storage, there is a significant growth delay for cells stored at either temperature. Cells stored at 4 °C did eventually approach full confluence, but those stored at −6 °C only reached 20% – 30% confluence after ten days.

Images of the cells that were stored for six days show that one day after plating, far fewer cells stored at −6 °C showed the spindle-like morphology of control cells, or even when compared to cells stored at 4 °C (Figure 7). Instead, the majority appear spherical. Additionally, the control cells show evidence of cell division as numerous cells clumps are visible, and even a small number of the cells that were stored at 4 °C appear to have divided. By day four, the number of cells visible in the −6 °C samples decreased, and a lower density of spindle-shaped cells remained. It is apparent by day seven that these cells are beginning to divide and form clumps, which is even more evident by day ten. However, these clusters are well-spaced, which indicates that only a small fraction of the cells that were originally plated have retained the ability to grow and divide. In contrast, those stored at 4 °C show a higher density of proliferative clusters at day seven, again suggesting that SZ-SCS is more deleterious to RPTEC/TERT1 cells than storage at 4 °C. As observed in the other assays, the addition of *Tm*AFP to CrS solution had no effect on the results.

**Figure 7.**
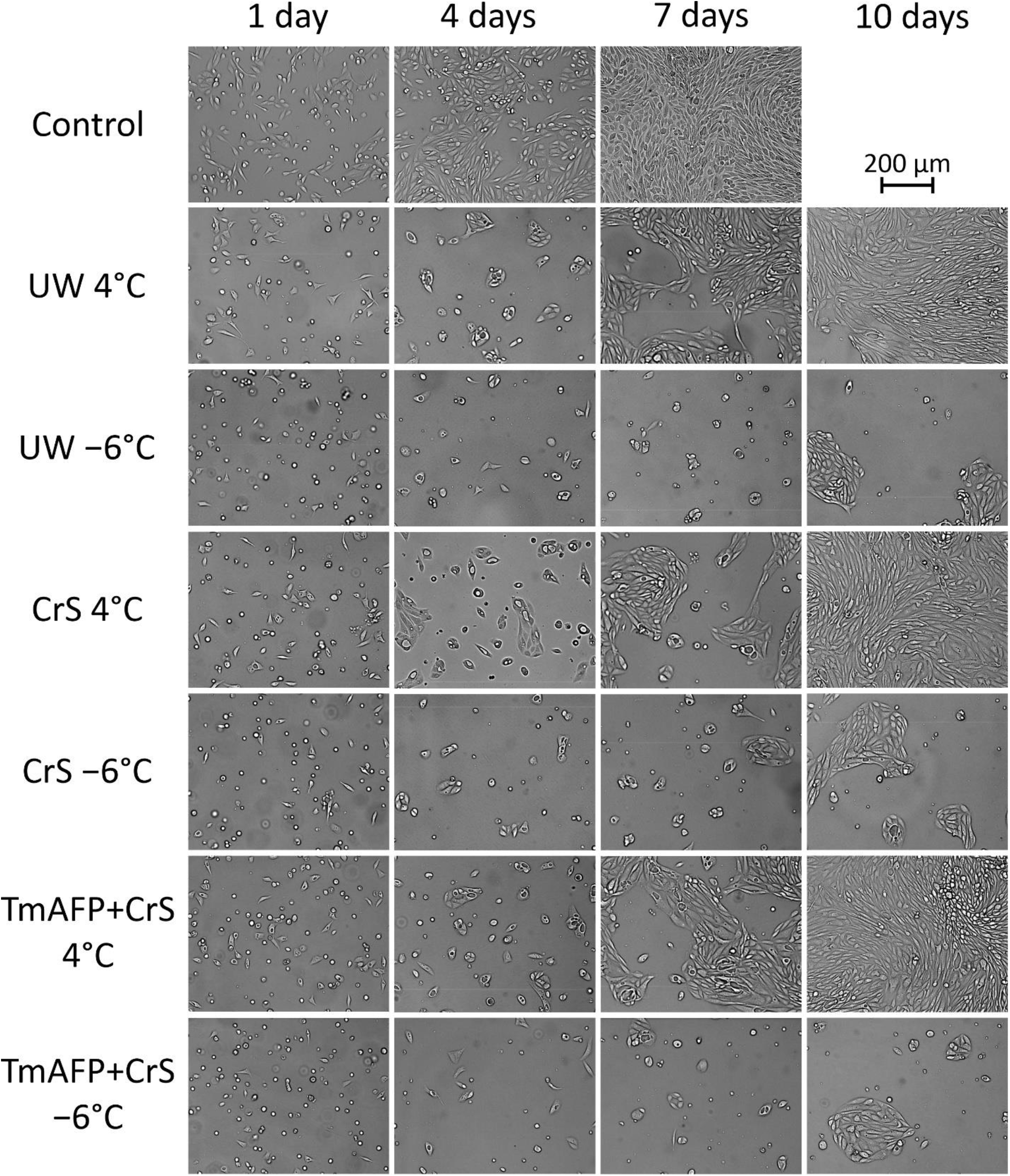
Representative images of a subset of the samples used to generate Fig. 6. All samples (except controls) were stored at the indicated temperatures for six days prior to plating. Contrast enhancement was applied to improve the visibility of the cells on the images.

## Discussion

The challenge of short organ storage times that limit the application of tissue matching has resulted in reduced success with transplants and translated to the loss of human life. Machine perfusion may provide a modest improvement in the transplant of kidneys and hearts ^2,26^, but long-term survival rates are not significantly different ^27^. Therefore, a less technically demanding alternative, such as or SZ-SCS, has been proposed as a possible solution to extend storage times and increase transplant success. Thus, we investigated subzero protocols to compare storage times of model human kidney cells stored at one, three and six days at −6 °C. As human organs are larger than those of rats and other small mammals frequently used in exploratory transplant studies, they are more difficult to maintain at subzero temperatures without freezing ^3^. Therefore, the ability of modest concentrations of the highly active *Tm*AFP to prevent ice formation at −6 °C, was examined. Even when challenged with the most potent biological ice nucleator known (*P. syringae* bacterial membranes with INP ^28^), *Tm*AFP was largely effective, suggesting it would stop organs with less potent nucleators from freezing.

Kidney failure, after injury or transplantation, often arises due to damage to the proximal tubule epithelial cells as they are particularly sensitive to ischemic-reperfusion injury, with necrosis, apoptosis and inflammatory processes playing an important role ^16-18,29^. The human RPTEC/TERT1 cells ^19^ used in this study are derived from kidney cells that undergo apoptosis in response to SCS, so they are a useful proxy with which to assess the potential effects of SZ-SCS on human kidneys. Unfortunately, the potential benefits of SZ-SCS, namely the 2-fold reduction in metabolic rate predicted by the 10-degree decrease in storage temperature ^5^ was eclipsed by damage associated with the low but non-freezing temperatures as evidenced by decreased cell viability, increased LDH release and apoptosis, reduced adhesion, and impaired proliferation.

Chilling injury is known to affect a number of cellular components during SCS. For example, mitochondria in aortic epithelial cells underwent fission during SCS ^30^, and microtubule disruption along with membrane dysfunction have been observed in the epithelial cells of kidneys following cold storage ^31^. Membranes are known to be sensitive to temperature changes as they undergo a phase transition from the liquid to the gel state as they are cooled ^32^, and this can vary between cell types, with lower transition temperatures correlating with reduced chilling injury ^33^. Additionally, some proteins can be denatured by exposure to low temperatures ^34,35^. Furthermore, significant levels of reactive oxygen species may be generated by mitochondria at −6 °C, since porcine kidneys stored at 4 °C generate over 50% of the hydrogen peroxide that they do at 37 °C, despite a 10-fold decrease in oxygen consumption ^36^. It is possible that any one of these effects, alone or in combination, is more detrimental to RPTEC/TERT1 at subzero temperatures than at 4 °C.

Chilling injury is not a barrier for all SZ-SCS protocols, as some studies using AFPs have shown varying degrees of success. For example, rat hearts stored at −1.3 °C in the presence of fish AFPs showed improved function relative to those stored in UW solution at 4 °C ^37,38^. Similarly, storage of rat kidneys with *Tm*AFP in CrS solution at −4.4 °C for three days showed promise, as several markers of tissue damage were equivalent to or lower than for kidneys stored in UW solution at 4 °C for only one day ^24^.

Additionally, high concentrations of *Tm*AFP were beneficial for SZ-SCS of pancreatic islet cells ^39^, but as these cells originated from a rat insuloma and have a reduction in chromosome number ^40^, they may be more resistant to chilling injury and apoptosis than RPTEC/TERT1 cells. Rat livers stored at −4 °C however, showed higher levels of LDH release and trypan blue uptake relative to those stored at 4 °C ^41^, similar to the results seen for RPTEC/TERT1 cells in this study.

There were no significant differences between cells stored in CrS solution and those in CrS solution with *Tm*AFP added. Therefore, *Tm*AFP was neither protective nor detrimental to RPTEC/TERT1 cells at a concentration of 0.1 mg/mL, indicating its utility in preventing freezing during SZ-SCS at temperatures as low as −6 °C. What is apparent however, is that SZ-SCS was detrimental to RPTEC/TERT1 cells that were derived from the epithelium of proximal tubules ^19^, a cell type known to be particularly sensitive to injury ^16,17^. Notably, while SZ-SCS appeared beneficial for rat kidney storage ^24^, those kidneys were not subsequently transplanted into recipients, so damage to the tubular epithelium, in particular, could not be assessed. In summary, the effectiveness of SZ-SCS varies depending on the tissue or cells type being preserved, and it may not be beneficial for storage of kidneys for transplantation, unless the specific mechanism that causes chilling injury to sensitive tubule epithelial cells can be identified and mitigated.

## Acknowledgements

In memory of Olga Kukal, co-founder of Cryostasis Ltd., whose tireless work in the field of cryopreservation led to the development of CrS solution (https://esc-sec.ca/wp-content/uploads/2021/02/2021_1-March-ESCBull-Final.pdf). This work was supported by Canadian Institutes of Health Research Foundation award (FRN 148422) to P.L.D., who holds the Canada Research Chair in Protein Engineering.

## References

1 Jing, L., Yao, L., Zhao, M., Peng, L. P. & Liu, M. Organ preservation: from the past to the future. Acta Pharmacol Sin 39, 845–857 (2018). https://doi.org:10.1038/aps.2017.182

2 Qin, G., Jernryd, V., Sjoberg, T., Steen, S. & Nilsson, J. Machine Perfusion for Human Heart Preservation: A Systematic Review. Transpl Int 35, 10258 (2022). https://doi.org:10.3389/ti.2022.10258

3 de Vries, R. J. et al. Supercooling extends preservation time of human livers. Nat Biotechnol 37, 1131–1136 (2019). https://doi.org:10.1038/s41587-019-0223-y

4 Peters-Sengers, H. et al. Impact of Cold Ischemia Time on Outcomes of Deceased Donor Kidney Transplantation: An Analysis of a National Registry. Transplant Direct 5, e448 (2019). https://doi.org:10.1097/TXD.0000000000000888

5 William, N. & Acker, J. P. High sub-zero organ preservation: A paradigm of nature-inspired strategies. Cryobiology 102, 15–26 (2021). https://doi.org:10.1016/j.cryobiol.2021.04.002

6 Berendsen, T. A. et al. Supercooling enables long-term transplantation survival following 4 days of liver preservation. Nat Med 20, 790–793 (2014). https://doi.org:10.1038/nm.3588

7 Rauen, U. & de Groot, H. Mammalian cell injury induced by hypothermia-the emerging role for reactive oxygen species. Biol Chem 383, 477–488 (2002). https://doi.org:10.1515/BC.2002.050

8 Best, B. P. Cryoprotectant Toxicity: Facts, Issues, and Questions. Rejuvenation Res 18, 422–436 (2015). https://doi.org:10.1089/rej.2014.1656

9 Pertaya, N. et al. Fluorescence microscopy evidence for quasi-permanent attachment of antifreeze proteins to ice surfaces. Biophys J 92, 3663–3673 (2007). https://doi.org:10.1529/biophysj.106.096297

10 Knight, C. A. Structural biology. Adding to the antifreeze agenda. Nature 406, 249, 251 (2000). https://doi.org:10.1038/35018671

11 Braslavsky, I. & Drori, R. LabVIEW-operated novel nanoliter osmometer for ice binding protein investigations. J Vis Exp, e4189 (2013). https://doi.org:10.3791/4189

12 Eskandari, A., Leow, T. C., Rahman, M. B. A. & Oslan, S. N. Antifreeze Proteins and Their Practical Utilization in Industry, Medicine, and Agriculture. Biomolecules 10 (2020). https://doi.org:10.3390/biom10121649

13 Bar Dolev, M., Braslavsky, I. & Davies, P. L. Ice-Binding Proteins and Their Function. Annu Rev Biochem 85, 515–542 (2016). https://doi.org:10.1146/annurev-biochem-060815-014546

14 Wang, L. & Duman, J. G. Antifreeze proteins of the beetle Dendroides canadensis enhance one another’s activities. Biochemistry 44, 10305–10312 (2005). https://doi.org:10.1021/bi050728y

15 Marshall, C. J., Basu, K. & Davies, P. L. Ice-shell purification of ice-binding proteins. Cryobiology 72, 258–263 (2016). https://doi.org:10.1016/j.cryobiol.2016.03.009

16 Cosio, F. G. et al. Kidney allograft fibrosis and atrophy early after living donor transplantation. Am J Transplant 5, 1130–1136 (2005). https://doi.org:10.1111/j.1600-6143.2005.00811.x

17 Schelling, J. R. Tubular atrophy in the pathogenesis of chronic kidney disease progression. Pediatr Nephrol 31, 693–706 (2016). https://doi.org:10.1007/s00467-015-3169-4

18 Grgic, I. et al. Targeted proximal tubule injury triggers interstitial fibrosis and glomerulosclerosis. Kidney Int 82, 172–183 (2012). https://doi.org:10.1038/ki.2012.20

19 Wieser, M. et al. hTERT alone immortalizes epithelial cells of renal proximal tubules without changing their functional characteristics. Am J Physiol Renal Physiol 295, F1365–1375 (2008). https://doi.org:10.1152/ajprenal.90405.2008

20 Tomalty, H. E., Graham, L. A., Eves, R., Gruneberg, A. K. & Davies, P. L. Laboratory-Scale Isolation of Insect Antifreeze Protein for Cryobiology. Biomolecules 9 (2019). https://doi.org:10.3390/biom9050180

21 Schneider, C. A., Rasband, W. S. & Eliceiri, K. W. NIH Image to ImageJ: 25 years of image analysis. Nat Methods 9, 671–675 (2012). https://doi.org:10.1038/nmeth.2089

22 Jaccard, N. et al. Automated method for the rapid and precise estimation of adherent cell culture characteristics from phase contrast microscopy images. Biotechnol Bioeng 111, 504–517 (2014). https://doi.org:10.1002/bit.25115

23 Schindelin, J. et al. Fiji: an open-source platform for biological-image analysis. Nat Methods 9, 676–682 (2012). https://doi.org:10.1038/nmeth.2019

24 Tomalty, H. E. et al. Kidney preservation at subzero temperatures using a novel storage solution and insect ice-binding proteins. Cryo Letters 38, 100–107 (2017).

25 Budke, C. & Koop, T. BINARY: an optical freezing array for assessing temperature and time dependence of heterogeneous ice nucleation. Atmos. Meas. Tech. 8, 689–703 (2015). https://doi.org:10.5194/amt-8-689-2015

26 Hosgood, S. A., Brown, R. J. & Nicholson, M. L. Advances in Kidney Preservation Techniques and Their Application in Clinical Practice. Transplantation 105, e202–e214 (2021). https://doi.org:10.1097/TP.0000000000003679

27 Kruszyna, T. & Richter, P. Hypothermic Machine Perfusion of Kidneys Compensates for Extended Storage Time: A Single Intervention With a Significant Impact. Transplant Proc 53, 1085–1090 (2021). https://doi.org:10.1016/j.transproceed.2021.01.022

28 Schwidetzky, R. et al. Membranes Are Decisive for Maximum Freezing Efficiency of Bacterial Ice Nucleators. J Phys Chem Lett 12, 10783–10787 (2021). https://doi.org:10.1021/acs.jpclett.1c03118

29 Bonventre, J. V. & Yang, L. Cellular pathophysiology of ischemic acute kidney injury. J Clin Invest 121, 4210–4221 (2011). https://doi.org:10.1172/JCI45161

30 Quiring, L. et al. Characterisation of cold-induced mitochondrial fission in porcine aortic endothelial cells. Mol Med 28, 13 (2022). https://doi.org:10.1186/s10020-021-00430-z

31 Breton, S. & Brown, D. Cold-induced microtubule disruption and relocalization of membrane proteins in kidney epithelial cells. J Am Soc Nephrol 9, 155–166 (1998). https://doi.org:10.1681/ASN.V92155

32 Arav, A. et al. Phase transition temperature and chilling sensitivity of bovine oocytes. Cryobiology 33, 589–599 (1996). https://doi.org:10.1006/cryo.1996.0062

33 Ghetler, Y., Yavin, S., Shalgi, R. & Arav, A. The effect of chilling on membrane lipid phase transition in human oocytes and zygotes. Hum Reprod 20, 3385–3389 (2005). https://doi.org:10.1093/humrep/dei236

34 Yan, R., Rios, P. D., Pastore, A. & Temussi, P. A. The cold denaturation of IscU highlights structure-function dualism in marginally stable proteins. Commun Chem 1 (2018). https://doi.org:10.1038/s42004-018-0015-1

35 Shan, B., McClendon, S., Rospigliosi, C., Eliezer, D. & Raleigh, D. P. The cold denatured state of the C-terminal domain of protein L9 is compact and contains both native and non-native structure. J Am Chem Soc 132, 4669–4677 (2010). https://doi.org:10.1021/ja908104s

36 Hendriks, K. D. W. et al. Renal temperature reduction progressively favors mitochondrial ROS production over respiration in hypothermic kidney preservation. J Transl Med 17, 265 (2019). https://doi.org:10.1186/s12967-019-2013-1

37 Amir, G. et al. Improved viability and reduced apoptosis in sub-zero 21-hour preservation of transplanted rat hearts using anti-freeze proteins. J Heart Lung Transplant 24, 1915–1929 (2005). https://doi.org:10.1016/j.healun.2004.11.003

38 Amir, G. et al. Prolonged 24-hour subzero preservation of heterotopically transplanted rat hearts using antifreeze proteins derived from arctic fish. Ann Thorac Surg 77, 1648–1655 (2004). https://doi.org:10.1016/j.athoracsur.2003.04.004

39 Yamauchi, A. et al. Subzero Nonfreezing Hypothermia with Insect Antifreeze Protein Dramatically Improves Survival Rate of Mammalian Cells. Int J Mol Sci 22 (2021). https://doi.org:10.3390/ijms222312680

40 Gazdar, A. F. et al. Continuous, clonal, insulin- and somatostatin-secreting cell lines established from a transplantable rat islet cell tumor. Proc Natl Acad Sci U S A 77, 3519–3523 (1980). https://doi.org:10.1073/pnas.77.6.3519

41 Soltys, K. A., Batta, A. K. & Koneru, B. Successful nonfreezing, subzero preservation of rat liver with 2,3-butanediol and type I antifreeze protein. J Surg Res 96, 30–34 (2001). https://doi.org:10.1006/jsre.2000.6053

42 Lane, L. B. Freezing Points of Glycerol and Its Aqueous Solutions. Industrial & Engineering Chemistry 17, 924–924 (1925). https://doi.org:10.1021/ie50189a017

43 Shipway, A. & Shipway, S. Depression of freezing point due to a substance in solution, <https://www.calctool.org/CALC/chem/substance/fp_depression> (2008).

44 Arai, T., Nishimiya, Y., Ohyama, Y., Kondo, H. & Tsuda, S. Calcium-Binding Generates the Semi- Clathrate Waters on a Type II Antifreeze Protein to Adsorb onto an Ice Crystal Surface. Biomolecules 9 (2019). https://doi.org:10.3390/biom9050162

45 Sonnichsen, F. D., Sykes, B. D., Chao, H. & Davies, P. L. The nonhelical structure of antifreeze protein type III. Science 259, 1154–1157 (1993). https://doi.org:10.1126/science.8438165

