## Supplementary Figures 1 and 2 for "Chill injury in human kidney tubule cells after subzero storage is not mitigated by antifreeze protein addition"

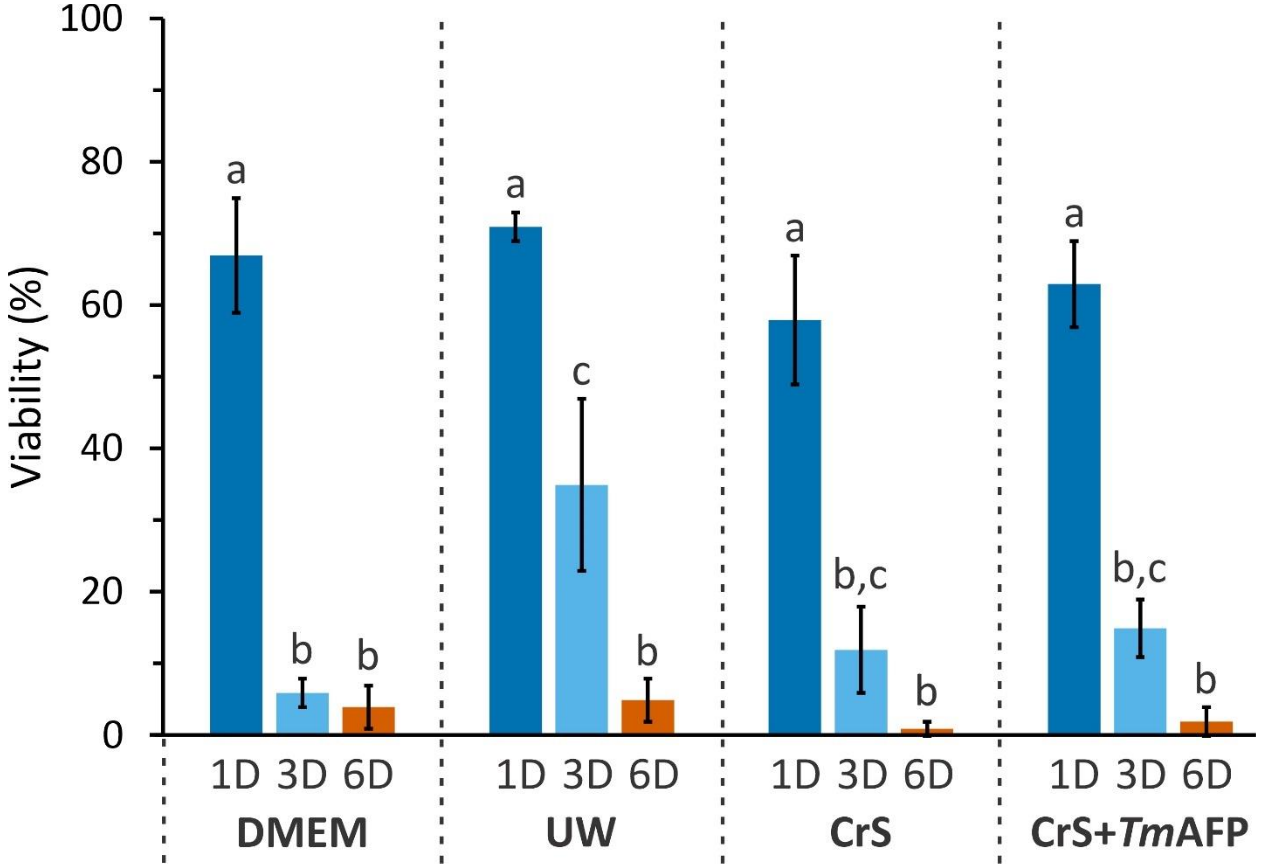


**Supplementary Figure 1.** Percentage of cells that survived storage under oil at 37 °C for one (dark blue), three (sky blue), or six days (reddish orange) in nutritive medium (DMEM) or storage medium (UW, CrS, CrS + AFP). The error bars represent the standard deviation from triplicates. Statistics are as described in Fig. 2 except that a two-way ANOVA was done.

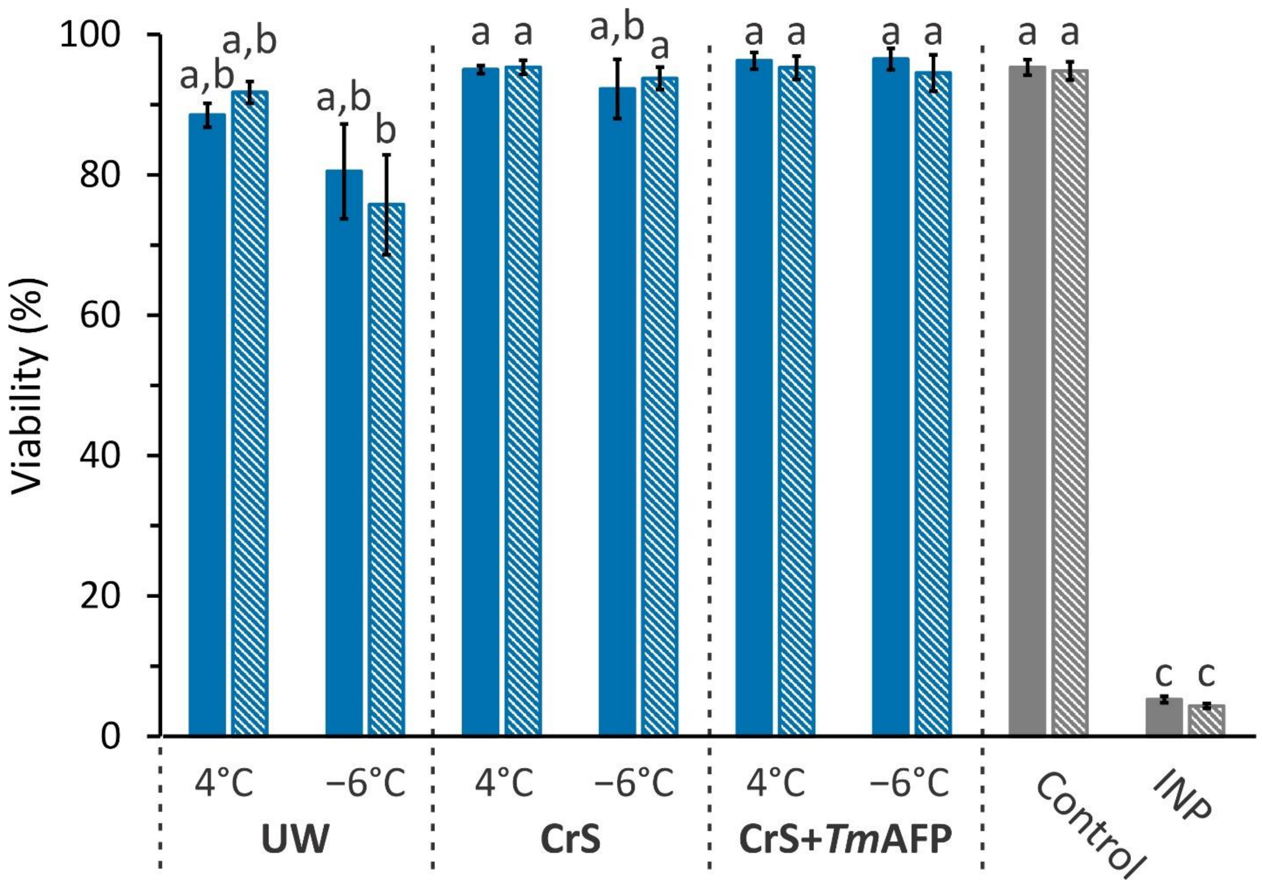


**Supplementary Figure 2.** Viability of RPTEC/TERT1 cells after conventional SCS (4 °C) or SZ-SCS (−6 °C) for one day in three different storage solutions (blue bars). Actively growing cells (control) and cells frozen in CrS at -6 °C by addition of ice nucleating protein (INP) are in grey. The recovery time at 37 °C was either 0 h (solid colours) or 20 h (hatched colours). The standard deviation of five replicates is shown. Statistics are as described in Fig. 2.
